# Phospholipase Dα1 acts as a negative regulator of high Mg^2+^-induced leaf senescence in Arabidopsis

**DOI:** 10.1101/2021.08.24.457483

**Authors:** Daniela Kocourková, Kristýna Kroumanová, Tereza Podmanická, Michal Daněk, Jan Martinec

**Affiliations:** Institute of Experimental Botany of the Czech Academy of Sciences, CZ-16502, Prague, Czech Republic

**Keywords:** *Arabidopsis thaliana*, magnesium homeostasis, phospholipase D, leaf senescence, starch, proline, abscisic acid, jasmonic acid, cytokinins

## Abstract

Magnesium is a macronutrient involved in essential cellular processes. Its deficiency or excess is a stress factor for plants, seriously affecting their growth and development and therefore, its accurate regulation is essential. Recently, we discovered that phospholipase Dα1 (PLDα1) activity is vital in the stress response to high-magnesium conditions in Arabidopsis roots. This study shows that PLDα1 acts as a negative regulator of high-Mg^2+^-induced leaf senescence in Arabidopsis. The level of phosphatidic acid produced by PLDα1 and the amount of PLDα1 in the leaves increase in plants treated with high Mg^2+^. A knockout mutant of PLDα1 (*plda1-1*), exhibits premature leaf senescence under high-Mg^2+^ conditions. In *pldα1-1* plants, higher accumulation of abscisic and jasmonic acid and impaired magnesium, potassium and phosphate homeostasis were observed under high-Mg^2+^ conditions. High Mg^2+^ also led to an increase of starch and proline content in Arabidopsis plants. While the starch content was higher in *plda1-1* plants, proline content was significantly lower in *plda1-1* compared with WT. Our results show that PLDα1 is essential for Arabidopsis plants to cope with the pleiotropic effects of high-Mg^2+^ stress and delay the leaf senescence.

## Introduction

Magnesium is a macronutrient involved in essential cellular processes such as photosynthesis, nucleic acid and protein synthesis, energy metabolism, etc. (Guo *et al*., 2016). Its deficiency or, on the contrary, its excess is a stress factor for plants, seriously affecting plant growth and development. Therefore, accurate regulation of intracellular magnesium level is essential. The knowledge of the mechanisms activated in Mg^2+^ deficiency is relatively good. The mechanisms associated with the regulation of cellular Mg^2+^ under high-Mg^2+^ conditions are less known. High concentrations of Mg^2+^ together with low concentrations of Ca^2+^ occur, for example, in serpentine soils. Recently, high-magnesium waters and soils have been considered as an emerging environmental and food security issues (Qadir *et al*., 2018). For non-adapted plants, high-Mg^2+^ conditions are strongly inhibitory to growth.

Magnesium is absorbed by plants in its ionic form from the soil. Plants growing in soils with high Mg^2+^ can mitigate Mg^2+^ toxicity by limiting internal Mg^2+^ accumulation and/or Mg^2+^ excretion from leaves. Sequestration of additional Mg^2+^ into the vacuole under high Mg^2+^ conditions appears to play a central role in tolerance to high Mg^2+^ (Hermans *et al*., 2013). The involvement of a network of calcineurin B-like calcium sensor proteins (CBL) CBL2/3, CBL-interacting protein kinases (CIPK) CIPK3/9/23/26, and sucrose nonfermenting-1-related protein kinase2 (SnRK2) SRK2D/E/I in the high-Mg^2+^ response of Arabidopsis has been shown (Chen *et al*., 2018; Mogami *et al*., 2015; Tang *et al*., 2015). Based on the altered sensitivity of the corresponding knock-out mutants to high-Mg^2+^ conditions, several other proteins were identified as participants in the high-Mg^2+^ response. Vacuolar-type H+-pyrophosphatase (AVP1) (Yang *et al*., 2018), magnesium transporter 6 (MGT6) (Yan *et al*., 2018), and *mid1*-complementing activity 1, 2 (MCA1/2) (Yamanaka *et al*., 2010) are required for tolerance to high Mg^2+^ because their knock-out mutants were hypersensitive to high-Mg^2+^ conditions. In contrast, knock-out mutants of cation exchanger 1 (CAX1) (Bradshaw, 2005; Cheng *et al*., 2003) and nucleoredoxin 1 (NRX1) (Niu *et al*., 2018) were more resistant to high Mg^2+^. Interestingly, MCA1/2, CAX1 and NRX1 are involved in the regulation of cytosolic Ca^2+^ concentration, suggesting a link between calcium homeostasis and high Mg^2+^ tolerance. In addition, the involvement of ABA signalling in response to high magnesium conditions has been demonstrated. An increase in ABA content and expression of ABA biosynthetic genes was reported under high magnesium conditions (Guo *et al*., 2014; Visscher *et al*., 2010). Moreover, the ABA - insensitive mutant *abi1-1* was less sensitive to high magnesium treatment than WT (Guo *et al*., 2014).

Recently, we discovered that phospholipase Dα1 activity is vital in the stress response to high-magnesium conditions in Arabidopsis. The T-DNA insertion mutant *pldα1* was hypersensitive to elevated magnesium levels and showed reduced primary root length and fresh weight. PLDα1 activity increases rapidly following high-Mg^2+^ treatment. Moreover, high-Mg^2+^ treatment was shown to disrupt K^+^ homeostasis.

Plant phospholipases D (PLD) cleave common phospholipids such as phosphatidylcholine releasing phosphatidic acid (PA) and free head group. PLDα1, the most abundant PLD member in Arabidopsis, has been reported to play a role in stress responses such as plant-microbe interaction, wounding, freezing, dehydration, and salinity (Hong *et al*., 2016; Ruelland *et al*., 2015; Wang *et al*., 2014). The PA apparently serves as a key signalling molecule in the above responses (Pokotylo *et al*., 2018).

Leaf senescence is a normal manifestation of plant ageing and represents the final stage of its development. There is also senescence induced by environmental stresses such as drought, cold, heat, low light, and pathogen attack (Sade *et al*., 2018; Zhang and Zhou, 2013), nutrient deficiencies such as nitrogen (Meng *et al*., 2016), potassium (Cao *et al*., 2006; Li *et al*., 2012; Wang *et al*., 2012), or magnesium (Tanoi and Kobayashi, 2015). Not only deficiency but also excess of nutrients leads to premature senescence of leaves. Exposure of sunflower plants to elevated K^+^ concentration resulted in premature leaf senescence (Santos, 2001). Ionic imbalance caused by salt stress also causes premature leaf senescence. The regulatory role of ROS (Allu *et al*., 2014) and transcription factor ANAC092 (Balazadeh *et al*., 2010) was revealed here. Interestingly, both ROS and ANAC092 are also involved in the regulation of developmental senescence. Thus, there is an overlap between stress-induced senescence and developmental senescence. In addition to ROS and specific transcription factors, the phytohormones jasmonic acid, ABA and cytokinins also play important roles in senescence processes.

This study shows that high external magnesium concentration triggers leaf senescence in Arabidopsis. Moreover, the knockout mutant of PLDα1 exhibits premature leaf senescence under high-Mg^2+^ conditions compared with WT. Under high-Mg^2+^ conditions, we also observed impaired ion homeostasis of *pldα1*. Furthermore, hormone, starch, and proline accumulation were altered in *pldα1* plants senescing under high Mg^2+^. From these results, we conclude that PLDα1 functions as a negative regulator of of high-Mg^2+^ induced leaf senescence.

## Materials and methods

### Plant materials

*Arabidopsis thaliana* Col-0 was used in the study. Knockout line *pldα1-1* (SALK_067533) was obtained from the NASC. Complemented lines *pldα1-1*Com1 and *pldα1-1*Com2 were described previously (Kocourková *et al*., 2020).

### Plant cultivation

Phenotypic experiments were performed on agar plates and in hydroponics. On vertical agar plates, plants were grown for 10 days on ½ MS, 1% agar (Sigma) and then transplanted into either control plates (½ MS (Duchefa), 1% agar) or high-Mg^2+^ plates (½ MS, 15 mM MgCl_2_, 1 % agar (Sigma)) and grown for another 7 days. Plates were kept in a growth chamber at 22°C during the day, 21°C at night, under long day (16 h of light) conditions at 100 μmol m^-2^ s^−1^ of light. Hydroponic plant cultivation was described in Kocourkova *et al*. (2020). 24-day-old hydroponically cultivated plants were treated with either ½ Hoagland’s solution (control) or ½ Hoagland’s solution with 15 mM MgSO_4_ added. The plants were grown in a growth chamber at 22 °C during the day (light intensity of 100 μmol m^−2^ s^−1^) and 21 °C at night in a 10-hour day/14-hour night mode.

### PLDα1 activity

To determine *in vitro* activity, extracts from a mixed leaf sample from the 3^rd^, 4^th^, 5^th^ and 6^th^ oldest true leaves of the plant (= mixed leaf sample) were prepared from hydroponically grown plants treated with 0 or 15 mM MgSO_4_ for 1, 2, 3, 4, 7 and 10 days. The leaves were frozen in liquid nitrogen. Samples were homogenized and buffer (per 1 mg sample 5µl buffer) consisting of 0.4 M sucrose, 0.1 M MgCl_2_, 0.1 M KCl, 50 mM HEPES-NaOH pH 7.5, Complete protease inhibitor coctail (Roche) and Pierce Phosphatase Inhibitor Mini Tablets (Thermo Fisher Scientific) was added to the homogenized samples. The samples were centrifuged for 10 min at 6010 g at 4 °C. The supernatant was transferred into a new tube and the samples were centrifuged for 90 min at 27400 g at 4 °C. The supernatant was collected and the protein concentration was measured using a Coomassie Plus Protein Assay (Thermo Scientific).

The enzymatic reaction 100µl contained 15 µl of sample (1µg /µl), 50 mM MES (pH 6.5, NaOH), 20 mM CaCl_2_ and 25 µl of substrate solution. The substrate solution contained 4 μM fluorescent PC (BODIPY-PC, Invitrogen™ by Thermo Fisher Scientific), 25 μM 1,2-dipalmitoyl-*sn*-glycerol-3-phosphocholine (Avanti Polar Lipids), 0.015% sodium deoxycholate and 50 mM MES buffer (pH 6.5). The substrate solution was incubated at room temperature for 30 minutes and then sonicated for 10 minutes. The reaction was started by adding the substrate and run for 30 min at 25 °C with shaking at 500 rpm. Lipids were extracted according to (Krckova *et al*., 2018). The lipids were separated first by the mobile phase methanol/chloroform/water/acetic acid (21/15/4/0.8) and after drying by the mobile phase chloroform/methanol/water (26/9/1). The plates were laser-scanned using Sapphire™ Biomolecular Imager (Azzure Biosystems) and evaluated using Azure Spot 2.2 software. The phosphatidic acid standard was prepared using commercial phospholipase D (Sigma Aldrich) (Pejchar *et al*., 2010).

### Western blot analysis

Western blot analysis was performed as described previously (Kocourková *et al*., 2020) with minor changes. Protein extracts were prepared as described above for TLC analysis. Proteins were separated on 10% SDS PAGE and transferred by wet blot overnight on a nitrocellulose membrane. PLDα1 protein was detected with anti-PLDα1/2 antibody (Agrisera) diluted 1: 2000 in 3% low fat milk in TBS-T. Goat anti-rabbit (Bethyl) in 5% low fat milk was used as a secondary antibody. Precision plus protein dual color standard (Biorad) was used and the position of the bands after blot transfer was marked on the membrane with a Western blot marker pen (Abcam). To control protein transfer, the membrane was stained with Novex reversible membrane protein stain (Invitrogen) according to the manufacturer’s instructions.

### Chlorophyll content

Samples were frozen in liquid nitrogen and homogenized. Chlorophyll was extracted into ethanol. Samples with ethanol were heated to 65 °C, left overnight at 4 °C and centrifuged (10 000 g, 10 min). The absorbance of the extracts was measured at 649 nm and at 665 nm and the chlorophyll content was calculated according to (Ritchie, 2006) and expressed as mg per g fresh weight.

### Gene transcription analysis

Gene transcriptions were measured either in whole aboveground parts of plants grown on agar treated with 0 or 15 mM MgCl_2_ for 7 days or in the mixed leave sample of plants grown hydroponically treated with 0 or 15 mM MgSO_4_ for 3 days. Measurement of gene expression was done according to (Kocourková *et al*., 2020) with minor changes. Briefly, RNA was isolated using the Spectrum Plant Total RNA Kit (Sigma-Aldrich) and genomic DNA removed using a Turbo DNA-free Kit (Applied Biosystems). Transcription was performed using the Transcriptor First Strand cDNA Synthesis Kit (Roche) with 0.5 µg RNA per reaction. Quantitative PCR was performed with a LightCycler 480 SYBR Green I Master Mix (Roche) on a LightCycler 480 System (Roche). The sequences of the primers used are listed in Table S2.

### Ion leakage

Rosettes of plants grown on agar and treated with 0 or 15 mM MgCl_2_ for 7 days were immersed in deionized water. Electrolyte leakage was measured with a COND 70 portable Conductivity Meter after 1 hour of incubation at room temperature. The samples were then autoclaved and the total conductivity of the extract was measured. The results were expressed as a proportion of the total conductivity in %.

### Measurement of nutrient content

Seedlings were grown for 10 d on half-strength MS media, after which they were transferred to agar plates with 0 or 15 mM MgCl_2_ for 7 d. Plates were kept in a growth chamber at 22°C during the day, 21°C at night, under long day (16 h of light) conditions at 100 μmol m^−2^ s^−1^. Samples (pooled plants, ∼100 mg dry weight) were digested with HNO_3_: HCl (6:1, v:v) and P, Mg^2+^, K^+^, and Ca^2+^ content was determined with inductively coupled plasma optical emission spectroscopy (Spectroblue, Spectro, Germany) analysis in the laboratory of Ekolab Žamberk, Czech Republic).

### Starch staining

For starch staining the 10-day-old plants grown on agar plates treated for 3 days with 0 or 15 mM MgCl_2_ and hydroponically grown 24-day-old plants treated for 3 days with 0 or 15 mM MgSO_4_ were used. Plants were collected at the end of the dark period. Chlorophyll was removed by immersion in 80% hot ethanol. The ethanol was changed until the rosettes were completely discolored. The rosettes were washed with water and then stained for 10 minutes with Lugol solution (Sigma) and washed for 1 hour in water at room temperature. The plants were then scanned on a Scanner Epson Perfection V800 Photo (Epson).

### Phytohormone analysis

Phytohormones were analyzed according to (Prerostova *et al*., 2021). Briefly, samples (20-45 mg FW leaves) were homogenized and extracted with 100 uL 50% acetonitrile solution. The extracts were centrifuged at 4° C and 30,000g. The supernatants were applied to SPE Oasis HLB 96-well column plates (10 mg/well; Waters, USA) and then eluted with 100 uL 50% acetonitrile. The pellets were then re-extracted. Phytohormones in each eluate were separated on Kinetex EVO C18 column (Phenomenex, USA). Hormone analysis was performed with a LC/MS system consisting of UHPLC 1290 Infinity II coupled to 6495 Triple Quadrupole Mass Spectrometer (Agilent, USA).

### Proline accumulation measurement

Proline content was measured in the mixed leaf sample of plants grown hydroponically treated with 0 or 15 mM MgSO_4_ for 3, 4 and 7 days with ninhydrin method (Bates *et al*., 1973). The samples were homogenized and proline was extracted into 3% sulfosalicylic acid (SSA, 5 µl/mg fresh weight). The samples were centrifuged (5 min at maximum speed) and supernatant was collected. The reaction mixture (180 µl) consisted of 30 µl sample, 96 µl glacial acetic acid, 24 µl 6 M orthophosphoric acid, 30 µl 3% SSA and 1.5 mg ninhydrin. The reaction was run for 1 hour at 96 °C. Then the samples were cooled on ice and 300 µl of toluene was added. The absorbance in the upper phase was measured at 520 nm.

## Results

### PLDα1 activity and amount of PLDα1 increase in leaves of Mg^2+^ treated plants

We had previously reported that increased Mg^2+^ concentration rapidly induces PLDα1 activity in Arabidopsis roots (Kocourkova *et al*., 2020). Here, we monitored PLDα activity after Mg^2+^ treatment in Arabidopsis leaves. 24-day-old hydroponically grown plants were treated with 15 mM MgSO_4_ for 1 to 10 days. Leaves from control and treated plants were harvested, homogenised, and the enzyme activity of PLDα was determined *in vitro*. PLDα activity increased throughout the observation period (1-10 days) compared with the control (Fig. 1A, B). Higher PLDα1 activity compared with the control was observed after two days of treatment with 15 mM MgSO_4_ (Fig. 1A, B) and increased 1.3-fold. The maximum activity was observed on the seventh day, when it increased almost 17-fold.

**Fig. 1.**
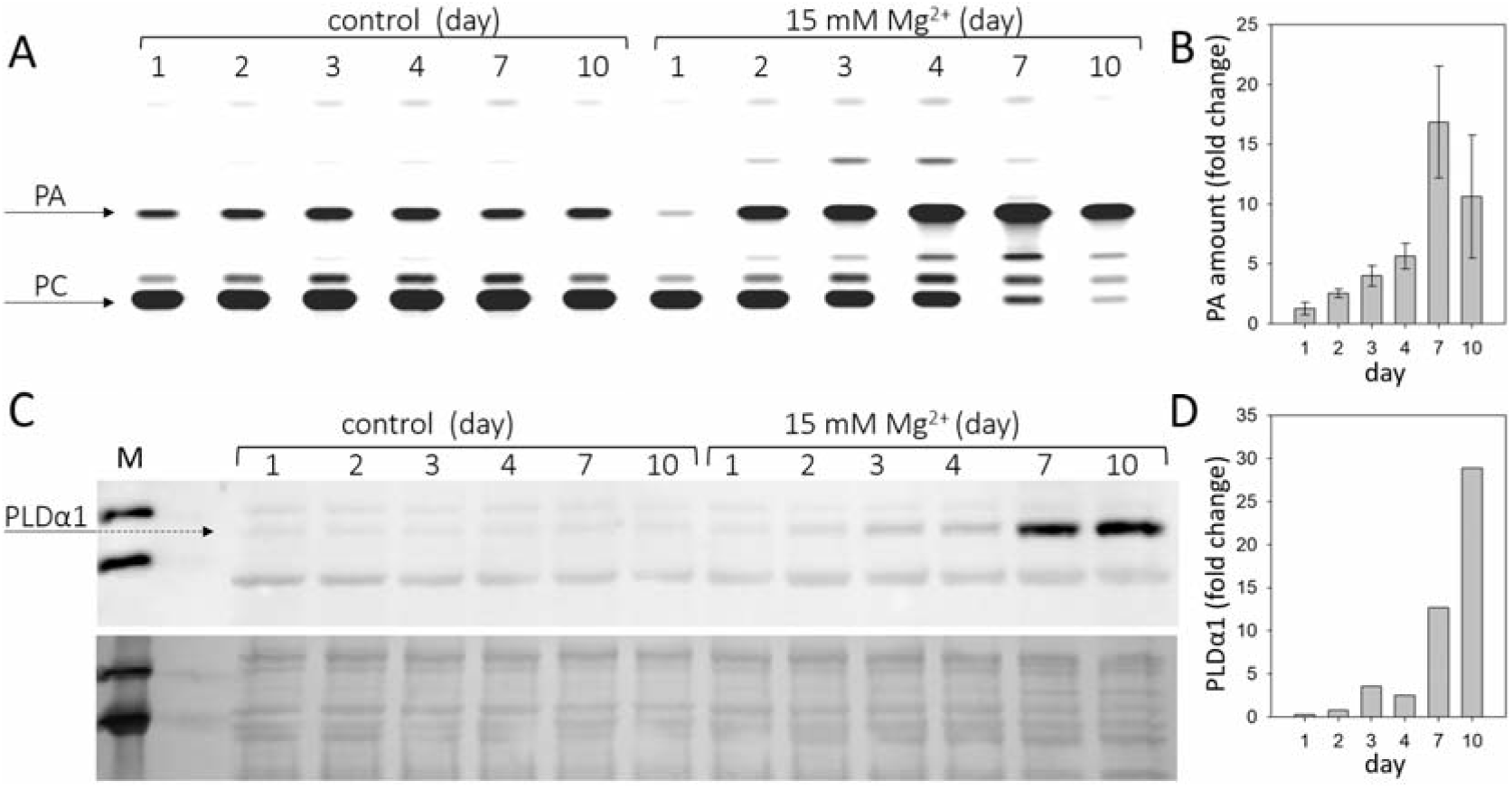
Phospholipase Dα1 activity and amount increases in response to high Mg^2+^ stress. 24-day-old hydroponically grown plants were treated with 15 mM MgSO_4_ and sampled after 1, 2, 3, 4, 7 and 10 days after the treatment. (A) Thin layer plate showing phosphatidic acid, product of PLDα1 activity. PLDα1 activity was measured in leave extracts from leaves 3-6 from plants treated with MgSO_4_. (B) Relative increase of PA with MgSO_4_ treatment over time. Values represent mean ± SE, n=3 biological experiments. (C) Western blot detection of PLDα1 in protein extracts from leaves. Each lane was run with 15 µg of protein, upper panel – western blot, lower panel – loading control - membrane stained with Novex reversible membrane protein stain. (D) Quantification of PLDα1 protein. The experiments were repeated three times with similar results. PA, phosphatidic acid, PC, phosphatidylcholine, M, molecular marker.

PLDs cleave common phospholipids such as phosphatidylcholine, releasing PA and the free head group, e.g. choline. PA is also the product of diacylglycerol kinase activity as well as the substrate for PA phosphatase, among other enzymes (Ruelland *et al*., 2015). Therefore, the PA level does not necessarily correlate with PLD activity. Moreover, there are several isoforms of PLD in Arabidopsis that differ in their biochemical properties (Kolesnikov *et al*., 2012). To measure PLD activity *in vitro*, we chose the optimal conditions for PLDα activity. PLDα activity was also determined in control and Mg^2+^-treated PLDα1 knockout plants (*pldα1-1*). In *pldα1-1*, no increase in PA level (PLDα activity) was observed under either control or high Mg^2+^ conditions (Fig. S1). Thus, the activity of the PLDα1 isoform is responsible for the observed increase in PA level. The increase in PLDα1 activity may be due to activation of PLDα1 or a higher level of PLDα1 protein, or both. The amount of PLDα1 in the leaves of control and treated plants was examined by western blot using the anti-PLDα1,2 antibody. The results clearly show that the level of PLDα1 increases after Mg^2+^ treatment (Fig. 1C, D). The difference between control and treated plants was detectable after three days of Mg^2+^ treatment.

Determination of the activity and level of PLDα1 was performed in samples consisting of mature third, fourth, fifth, and sixth leaves. Senescence symptoms were slightly visible in these leaves on the seventh day. However, the same trend, increased activity of PLDα1, was observed in the young leaves (7th -10th), which showed no visible signs of senescence (Fig. S2). Therefore, we hypothesise that changes in PLDα1 activity and content are not downstream of the manifestation of leaf senescence.

These results show that both PLDα1 activity and PLDα1 levels increase in Arabidopsis leaves after treatment with Mg^2+^.

### High magnesium induces premature leaf senescence in pldα1

We found (Kocourkova *et al*., (2020) that 12-day-old Arabidopsis seedlings of *pldα1* under high-Mg^2+^ conditions had shorter primary and lateral roots and lower fresh weight. Here, we noticed higher yellowing or yellow spots on *pldα1-1* leaves after Mg^2+^ treatment (15 Mm MgSO_4_) of 24-day-old plants (Fig. 2A, B). Also, the fresh weight of *pldα1-1* rosettes was less than half compared with WT, and the chlorophyll content of *pldα1-1* decreased by about 35% compared with WT (Fig. 2C, D). This indicates premature leaf senescence of *pldα1-1* plants. To further characterise the observed phenomenon, we additionally monitored high-Mg^2+^-induced senescence by determining the expression of senescence genes and measuring ion leakage as a marker of membrane damage. To verify that the observed premature senescence was exclusively related to PLDα1, we also included two *pldα1-1*complemented (*pldα1-1*Com1 and Com2) lines (Kocourková *et al*., 2020). As in the adult plants, the fresh weight of *pldα1-1* rosettes was significantly lower (36%) compared with the WT and complemented lines (Fig. 2F). Expressions of Senescence-Associated Genes 13 (SAG13, At2g29350) and the transcription factor ANAC092/NAC2/ORE1 (At5g39610) are commonly used as markers of senescence (Balazadeh *et al*., 2010; Bresson *et al*., 2018). Expressions of these genes were higher in all Mg^2+^-treated WT, *pldα1-1* and complemented plants in comparison with untreated controls. Hence, high-Mg^2+^ induced transcriptional changes accompanying leaf senescence in all studied genotypes. Moreover, in *pldα1-1* plants, *SAG13* and *ANAC092* expression was notably higher than in WT or complemented lines. Expression of *SAG13* was approximately 2000-fold higher in *pldα1-1* seedlings treated with high Mg^2+^ than in the untreated control, whereas for WT the increase was only 314-fold higher than in the untreated control (Fig. 2G). Similarly, the expression of *ANAC092* was increased 12-fold in *pldα1-1*, whereas it increased only 2-fold in WT (Fig. 2H). Membrane damage was estimated by measuring ion leakage. After Mg^2+^ treatment, ion leakage reached 7.7% in *pldα1-1*, while it was only 3% in WT.

**Fig. 2.**
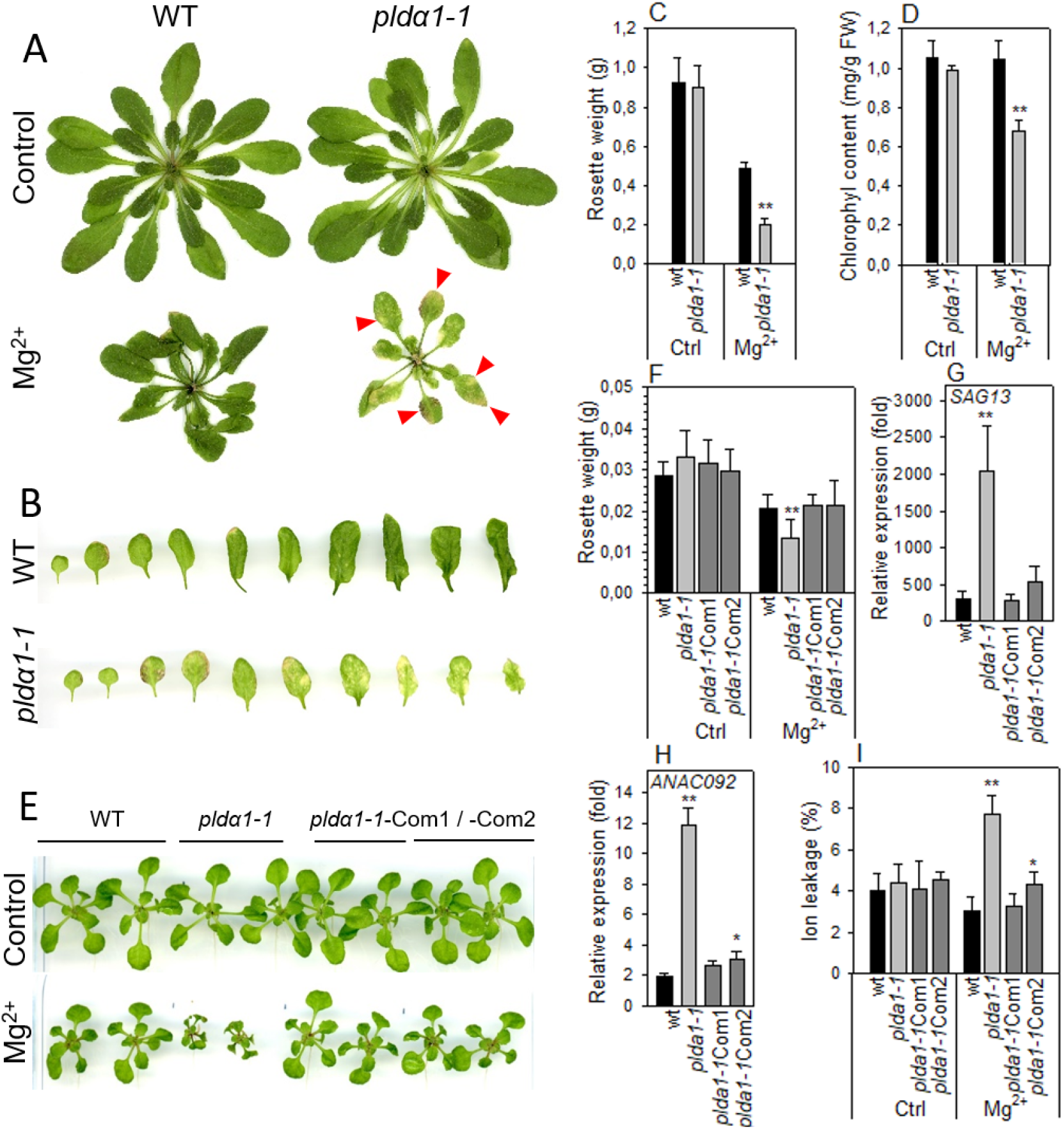
High Mg^2+^ leads to premature senescence in *pldα1-1*. (A) Phenotype of WT and *pldα1-1* grown hydroponically on high Mg^2+^, red arrows point to senescent parts of leaves. (B) Leaves of plants grown on high Mg^2+^. (C) Weight of rosettes. (D) Chlorophyll content, A-D - three-week-old hydroponically grown plants were treated with 15 mM MgSO_4_ and grown for another 16 days, values represent means ± SD, n=6. (E) Phenotype of WT, *pldα1-1, pldα1-1*Com1 and *pldα1-1*Com2 plants grown on agar plates on high Mg^2+^. (F) Weight of rosettes. (G), (H) Transcript level of *SAG13* and *ANAC092* in rosettes. Transcription was normalised to a reference gene *SAND* and the transcription of non-treated plants was set to one. Values represent means ± SE, n=12. (I) Ion leakage, values represent means ± SD, n=7. E-I - ten-day-old Arabidopsis seedlings were transferred on agar plates containing 15 mM MgCl_2_ and grown for another 7 days, Student’s T-test, asterisk indicate significant difference in comparison with WT * P<0.05; ** P<0.01.

These results demonstrate that high-Mg^2+^ treatment induces premature leaf senescence and that *pldα1-1* plants reveal significantly higher premature leaf senescence in comparison to WT. Premature senescence after high-Mg^2+^ treatment was observed in both ten-day-old seedling and three-week-old mature plants. Thus, we hypothesise that PLDα1 acts as negative regulator of high-Mg^2+^ induced leaf senescence.

### Levels of plant hormones are altered in high-Mg^2+^ conditions

Plant hormones are one of the key components involved in the processes of leaf senescence, influencing all stages, initiation, progression and terminal phase, of leaf senescence (Lim *et al*., 2007). Additionally, Guo *et al*. (2014) reported increase level of abscisic acid (ABA) in response of Arabidopsis Landsberg erecta to high-Mg^2+^ conditions. Hence, we measured range of phytohormones in WT and *pldα1-1* in control and high-Mg^2+^ conditions after two days of high-Mg^2+^ treatment (Table S1). Principal component analysis of all measured shoot phytohormones showed a clear separation on the PC1 axis of both control and high-Mg^2+^ conditions and genotypes (WT vs *pldα1-1*). There was also a separation on the PC2 axis between WT and *pldα1-1* genotype in high-Mg^2+^ conditions (Fig. 3A). These results demonstrate robust hormonal response to high-Mg^2+^ conditions in WT as well as involvement of PLDα1 in this hormonal response.

**Fig. 3.**
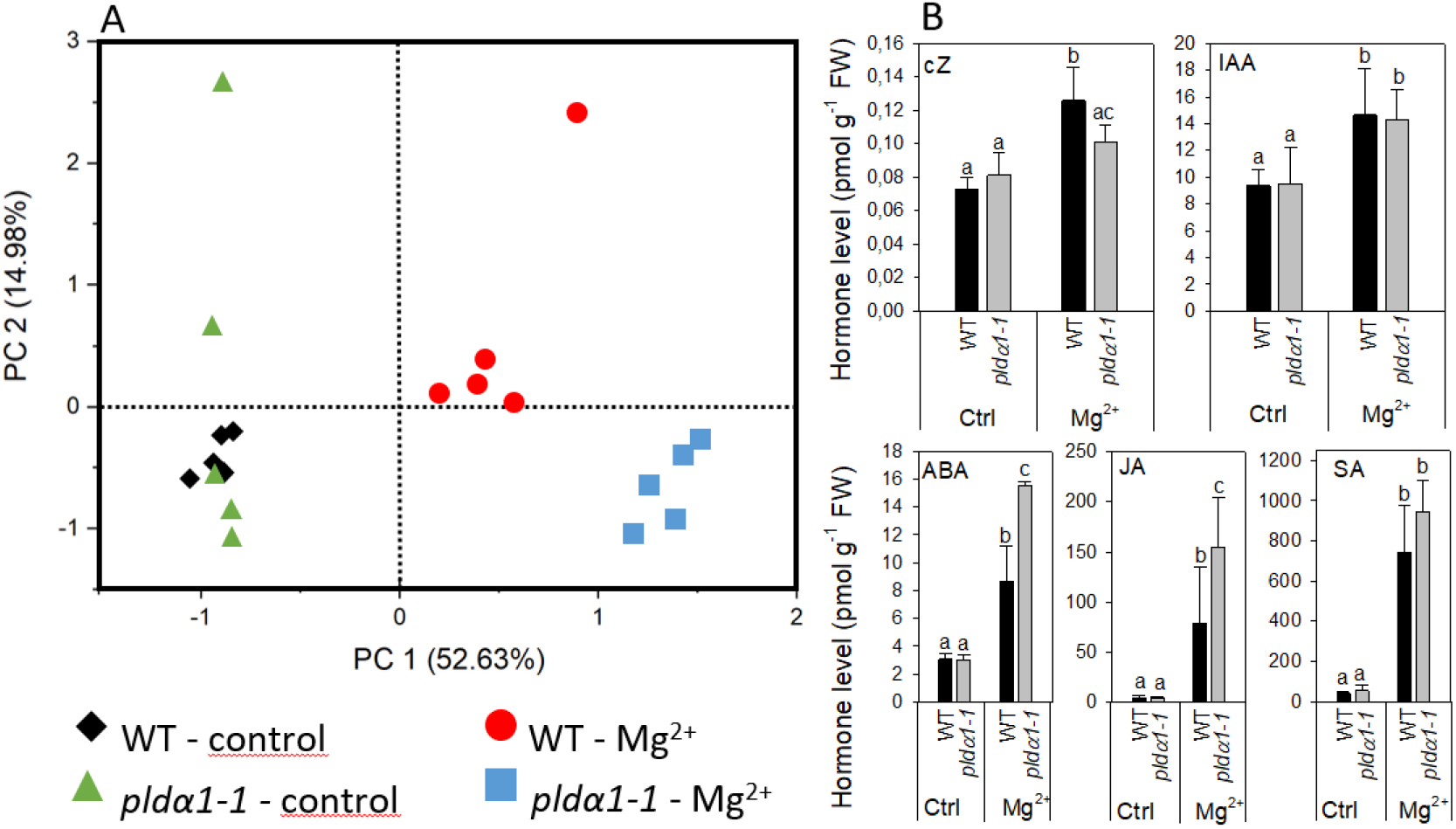
Levels of plant hormone are altered in high-Mg^2+^ conditions. (A) PCA analysis. (B) Level of cZ – cis-zeatin, IAA – indole-3-acetic acid, ABA – abscisic acid, JA – jasmonic acid and SA – salicylic acid. 24-day-old hydroponically grown plants were treated with 15 mM MgSO_4_ and grown for another 2 days. Leaves 3-6 were used for hormone analysis. Values represent means ± SD, n=5, letters above the bars indicate significant differences, one-way ANOVA with Tukey’s post hoc test, p<0.05.

In WT, the highly active cytokinin (CK) trans-zeatin (tZ) and its riboside (tZR) lowered after Mg^2+^ treatment. Also, the content of the precursor trans-zeatin riboside monophosphate (tZRMP) lowered in high Mg^2+^ treated shoots. On the opposite, the levels of the stress-related CKs cis-zeatin (cZ), its riboside (cZR), and phosphate (cZRMP) increased after high Mg^2+^ treatment in WT.

The high-Mg^2+^ treatment up-regulated the production of auxin indole-3-acetic acid (IAA) in WT plants. The level of IAA precursor, indole-3-acetamide (IAM) increased under the same conditions as well. Also, deactivation of production of IAA irreversible amino acid conjugate, IAA-glutamate significantly decreased after high Mg^2+^ treatment in WT.

ABA and its catabolites phaseic acid and 9-hydroxy-abscisic acid (9OH-ABA) elevated about three times in WT shoot of high-Mg^2+^ treated plants in comparison with non-treated plants. Jasmonic acid (JA) was greatly up-regulated (about 17 times) in high-Mg^2+^ conditions. Similarly, levels of JA precursor, cis-12-oxo-phytodienoic acid (cisOPDA) and JA metabolites, jasmonic acid methyl ester (JA-Me) and dinor-12-oxo-phytodienoic acid (dinorOPDA) significantly increased. Great increase of shoot salicylic acid (SA) level was detected in high-Mg^2+^ treated WT plants as well.

Under control conditions, no significant differences were found between WT and *pldα1-1* in the levels of all hormones measured, except for tZR. Under high-Mg^2+^ conditions, changes of some of the hormones differed between WT and *pldα1-1* (Fig. 3B, Table S1). Increase of SA was the same in WT and *pldα1-1*. Also, the increase of IAA was the same in WT and *pldα1-1*. However, higher increase was observed in the levels of both IAA precursor, IAM and IAA metabolite oxo-IAA-glucose ester (OxIAA-GE). Increase of JA (but not its precursor or metabolites) was significantly higher in *pldα1-1* in comparison with WT (Fig. 3A, Table S1). Increase of ABA as well as its catabolites phaseic acid and 9OH-ABA was more pronounced in *pldα1-1* than in WT. Interestingly, increase of cZ detected after Mg^2+^ treatment in WT was not observed in *pldα1-1*.

These results revealed that high-Mg^2+^ condition induce range of hormonal changes in both WT and *pldα1-1* plants. However, changes in ABA and JA levels observed after treatment of plants with high-Mg^2+^ were more pronounced in *pldα1-1* plants. Thus, it suggests that those hormonal changes are, at least partly, under the control of PLD*α*1. It means that the function of PLDα1 in regulation of hormonal changes after high-Mg^2+^ treatment is specific, as the observed difference between WT and *pldα1-1* did not affect all hormones that changed after high-Mg^2+^ treatment of plants but only ABA, JA and cis-zeatin.

### Ion homeostasis and levels of starch and proline are altered in pldα1-1 under high-Mg^2+^ conditions

In our previous work, an imbalance of K^+^ and Mg^2+^ was found in the seedlings of *pldα1-1* treated with high-Mg^2+^. They contained less Mg^2+^ and K^+^ under high-Mg^2+^ conditions (Kocourková *et al*., 2020). To reveal whether a similar ion imbalance also occurs in high-Mg^2+^-treated shoots, we measured Mg^2+^, Ca^2+^, K^+^ and P in WT and *pldα1-1* shoots under control and high-Mg^2+^ conditions.

After high-Mg^2+^ treatment (15 mM), Mg^2+^ content was increased approximately fivefold in WT leaves. However, *pldα1-1* showed significantly lower Mg^2+^ content than WT (Fig. 4A). Shoot K^+^ content was lower in high-Mg^2+^-treated plants, and *pldα1-1* plants contained even less K^+^ than WT (Fig. 4B). Ca^2+^ content was lower in high-Mg^2+^-treated plants, but WT and *pldα1-1* content did not differ (Fig. 4C). Furthermore, Niu *et al*. (2015) showed that the addition of phosphorus to high-Mg^2+^ media resulted in an increase in Arabidopsis root growth and, conversely, the addition of high-Mg^2+^ to low-P media worsened root growth. Based on these results, the authors speculated that the exacerbation of the effects of low P in the presence of high Mg^2+^ was due to the increase in the severity of P deficiency. Therefore, we also measured P content in WT and *pldα1-1* grown on high Mg^2+^ media. Remarkably, the phosphorus content in *pldα1-1* shoots under high-Mg^2+^ conditions was significantly lower than in WT (Fig. 4D).

**Fig. 4.**
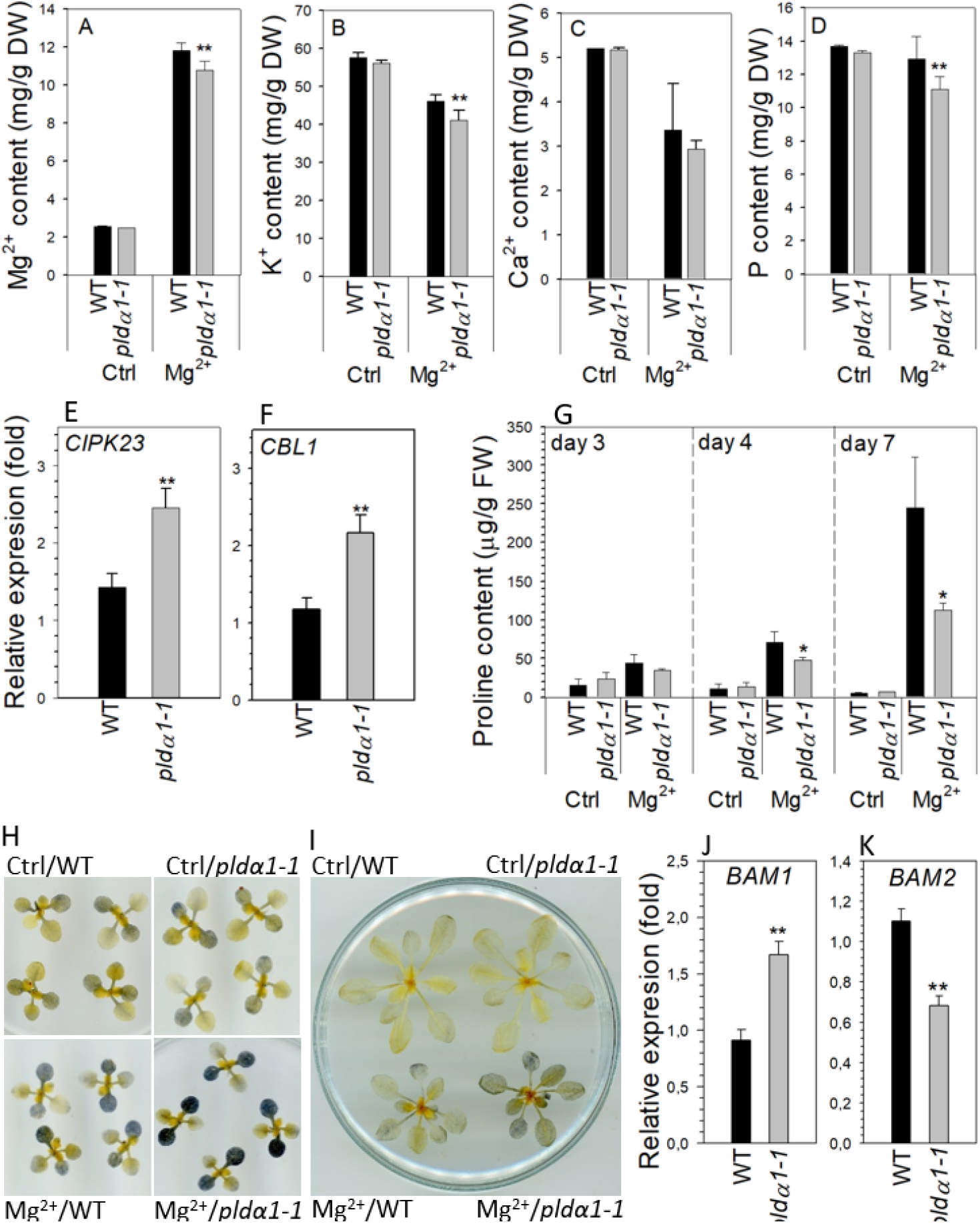
Nutrient, proline and starch content is altered in *pldα1-1* in high-Mg^2+^ conditions. (A), (B), (C), (D) Magnesium, potassium, calcium and phosphorus content in rosettes of WT and *pldα1-1* plants. Bars represents means ± SD, n=3 for control, n=8 for Mg treatment. (E), (F) Transcript level of *CIPK23* and *CBL1* in rosettes. Transcription was normalised to a reference gene *SAND* and the transcription of non-treated plants was set to one. Values represent means ± SE, n=12, A-F 10-day-old Arabidopsis seedlings were transferred on agar plates containing 15 mM MgSO_4_ for 7 days. (G) Proline content, 24-day-old hydroponically grown plants were treated with 15 mM MgSO_4_. Proline content was measured in leaves 3-6. (H) Lugol staining of starch in rosettes from control and high Mg^2+^ condition at the end of dark period. 10-day-old Arabidopsis seedlings were transferred on agar plates containing 15 mM MgSO_4_ for 3 days. (I) Lugol staining of starch in rosettes from control and high Mg^2+^ condition at the end of dark period, 24-day-old hydroponically grown plants were treated with 15 mM MgSO_4_ for 3 days. (J), (K) Transcript level of *BAM1* and *BAM2* in leaves 3-6, 24-day-old hydroponically grown plants were treated with 15 mM MgSO_4_ for 2 days, transcription was normalised to a reference gene *SAND* and the transcription of non-treated plants was set to one. Values represent means ± SE, n=12. Student’s T-test, asterisk indicate significant difference in comparison with WT * P<0.05; ** P<0.01.

These results were supported by determining the expression of genes known to be associated with ion homeostasis. CBL1 is involved in potassium as well as phosphate homeostasis, whereas CIPK23 is thought to be involved in both magnesium and potassium homeostasis (Gao *et al*., 2020; Ragel *et al*., 2015; Sánchez-Barrena *et al*., 2020; Tang *et al*., 2015). In WT leaves, the expression of these genes was slightly up-regulated under high-Mg^2+^ conditions (Fig. 4E, F). However, in *pldα1-1* leaves treated with high Mg^2+^, *CIPK23* and *CBL1* transcripts were significantly higher than in WT plants.

Proline is well known stress molecule involved mainly in responses to drought and salt stress. Increase of proline content was also reported as response to phosphate starvation in Arabidopsis (Aleksza *et al*., 2017). As we observed decrease of phosphate content in high-Mg^2+^ treated *pldα1-1* plants we monitor proline level in high-Mg^2+^ treated WT and *pldα1-1* leaves. Proline content substantially increased with increasing time of Mg^2+^ treatment and was tenfold higher in WT plants treated for seven days with high Mg^2+^ than in control plants. Interestingly, shoots of *pldα1-1* contained significantly lower level of proline after Mg^2+^ treatment in comparison with WT (Fig. 4G).

It has been reported that that high-Mg^2+^ treatment disturbs starch homeostasis (Guo 2014-citace) and that both potassium and phosphorus deficiency lead to accumulation of leaf starch (Hermans *et al*., 2006; Hu *et al*., 2017). We stained starch with Lugol’s solution in WT and *pldα1-1* seedlings and 24-day-old plants grown under control and high-Mg^2+^ conditions. At the end of dark period, there was clearly a higher starch accumulation in the shoot after Mg^2+^ treatment (Fig. 4H, I). Interestingly, higher starch accumulation was observed in *pldα1-1* compared with WT plants. Moreover, shoot expression of β-amylases *BAM1* and *BAM2* was impaired in *pldα1-1* compared with WT under high Mg^2+^ treatment (Fig. 4J, K).

These results demonstrate that Mg^2+^, K^+^, and P homeostasis, starch metabolism and proline accumulation are altered in *pldα1-1* shoots of Arabidopsis seedlings grown under high-Mg^2+^ conditions.

## Discussion

In our previous work (Kocourková *et al*., 2020) we found that *pldα1* plants have shorter roots under high Mg^2+^ conditions compared to WT. We also showed that PLDα1 activity contributes significantly to tolerance to high Mg^2+^. In this work, we focused on the shoots. Our original hypothesis was that PLDα1 activity is important mainly in roots, as they are exposed to high Mg^2+^ conditions and an increase in PLD activity is rapidly induced after high Mg^2+^ treatment (Kocourková *et al*., 2020). However, we found that PLDα1 activity in the aerial parts of WT Arabidopsis also increased after treatment of the plants with high level of Mg^2+^ ions. Using western blots, we also showed that the amount of PLDα1 increased in shoots treated with high Mg^2+^ and that *pldα1* plants exhibited premature leaf senescence under high Mg^2+^ conditions. Thus, PLDα1 appears to act as a negative regulator of senescence induced by high Mg^2+^.

### Magnesium-induced senescence

Leaf senescence is a highly coordinated process. In addition to age-dependent senescence, there is also stress-induced senescence caused by abiotic (drought, salt, high or low temperature, and nutrient imbalance) and biotic stresses (Guo *et al*., 2021; Sade *et al*., 2018). Leaf senescence is associated with membrane and chlorophyll degradation. Leaf yellowing and senescence have been reported to be induced by magnesium deficiency (for a review, see (Tanoi and Kobayashi, 2015). In addition, a decrease in chlorophyll content was observed by Yan *et al*. (2018) under conditions of Mg^2+^ imbalance (Mg deficiency and excess). We observed a greater decrease in leaf chlorophyll content and higher ion leakage in *pldα1* plants than in WT. Also, the transcript level of the senescence marker genes *SAG13* (Dhar *et al*., 2020) and *ANAC092* (John *et al*., 2001; Miller *et al*., 1999; Weaver *et al*., 1998) was significantly higher in *pldα1* plants, although the expression of both genes was also increased in WT plants in which no signs of senescence were yet evident. Since both genes are among the markers of the onset of senescence, it can be concluded that senescence processes are also initiated in WT upon high Mg^2+^ treatment. However, plants with dead PLDα1 tolerate the stress caused by high Mg^2+^ concentrations much worse than WT, leading to apparent premature leaf senescence.

### The high Mg^2+^ induced senescence-associated hormonal changes

Hormones play a critical role in regulating both development and stress-induced senescence. Cytokinins (CK), auxin and gibberellic acid (GA) delay leaf senescence, while abscisic acid (ABA), salicylic acid (SA), jasmonic acid (JA), ethylene and strigolactones (SL) promote leaf senescence (Guo *et al*., 2021; Lim *et al*., 2007). The overall hormonal changes we observed after high Mg^2+^ treatment of WT plants were in good agreement with the reported hormonal changes during leaf senescence. We found that after two days of high Mg^2+^ treatment, there was a decrease in active cytokinins, such as *tran*s-zeatin (tZ) and its riboside (tZR) which is in line with gradual decrease in cytokinin content observed during leaf senescence (Gan and Amasino, 1996; Singh *et al*., 1992). On the other hand, the content of the stress-related CKs *cis*-zeatin (cZ), its riboside (cZR), and phosphate (cZRMP) increased. An increase in cZ during natural senescence was reported in Arabidopsis and tobacco (Gajdošová *et al*., 2011; Uzelac *et al*., 2016). The level of cZ differed in WT and *pldα1* plants under high Mg^2+^ conditions. This suggests that the level of cZ is regulated by PLDα1 under high Mg^2+^ conditions. However, the level of cZ in more senescent *pldα1* plants is lower compared with WT. This is a counterintuitive finding, and further studies are required to clarify this phenomenon.

Auxin seems to play a complex role in senescence. On the one hand, its level and the level of auxin-related biosynthetic genes increase during natural leaf senescence (Lim *et al*., 2007). On the other hand, increased IAA level is associated with delayed dark-induced senescence (Kim *et al*., 2011). In our experiments, the level of free IAA increased significantly in WT and *pldα1* plants treated with high Mg^2+^, but we found no difference between genotypes, indicating that the observed phenotypic changes do not appear to be related to auxin content.

In our experiment, a significant increase in salicylic acid (SA) was observed after only two days of high Mg^2+^ treatment. Salicylic acid is a hormone well described in the mechanism of response to biotic stress, but it is also associated with responses to abiotic stress (Miura and Tada, 2014). Moreover, the level of SA increases progressively during leaf senescence (Breeze *et al*., 2011; Zhang *et al*., 2017), and SA plays a direct role in both the onset and progression of leaf senescence (Guo *et al*., 2021).

Abscisic acid (ABA) is a plant hormone whose level increases significantly after abiotic stresses such as drought and salt stress. During leaf senescence, the level of ABA increases, and exogenous application of ABA induces leaf senescence (Lim *et al*., 2007; Zhang *et al*., 2012a). We also observed a significant increase in ABA level after high Mg^2+^ treatment in WT and *pldα1* plants, while the increase of ABA was higher in *pldα1* plants than in WT plants. This is consistent with the higher senescence of *pldα1* induced by high Mg^2+^ content, which is also consistent with the observations of Guo *et al*. (2014). The authors found an increase in ABA content after long-term (14 d) exposure of Arabidopsis to high Mg^2+^. They also showed that ABA insensitive plants *abi1-1* were less sensitive to high Mg^2+^ treatment. In our experiments, a significant ABA response was observed after only 48 h of exposure. Moreover, transcriptome analysis of Arabidopsis roots treated with high Mg^2+^ revealed increased expression of 9-cis-epoxycarotenoid dioxygenase, an enzyme associated with the biosynthesis of ABA, after 45 min of high Mg^2+^ treatment (Visscher *et al*., 2010). All these results suggest that the increase in ABA content and subsequent ABA signalling are involved in the PLDα1-mediated early responses to high Mg^2+^ conditions.

Like SA, jasmonic acid (JA) is a hormone known primarily for its involvement in plant defence mechanisms against pathogens. However, it is also associated with many responses to abiotic stresses, such as salt, heat, or drought stress (Raza *et al*., 2020). It has been shown that JA content increases during both natural and induced leaf senescence, and external application of JA induces leaf senescence (He *et al*., 2002; Seltmann *et al*., 2010), so JA plays a positive role in leaf senescence. However, plants with impaired JA biosynthesis showed normal leaf senescence (Seltmann *et al*., 2010), suggesting that JA is not directly involved in the regulation of leaf senescence. In our experiments, a significant increase in JA was observed in both WT and *pldα1* plants after high Mg^2+^ treatment, and this increase was more pronounced in *pldα1* plants.

JA signalling has been shown to play a role in the biosynthesis of camalexin (Pangesti *et al*., 2016), a phytoalexin with a described role in the defence response to a variety of pathogens. Its biosynthesis is also induced by some abiotic treatments such as ROS-inducing compound acifluorfen (Zhao *et al*., 1998) or UV-B irradiation (Mert-Turk *et al*., 2003). Interestingly, camalexin content increased after high Mg^2+^ treatment in WT and even more (by 4.5-fold) in *pldα1* plants (Table S1). It is not clear what role camalexin might play in response to high Mg^2+^ treatment. However, higher camalexin levels in *pldα1* plants might be related to higher JA levels in *pldα1* plants treated with high Mg^2+^. In addition, camalexin biosynthesis is regulated by MPK6 kinase, which has been shown to be a PA binding protein (Yu *et al*., 2010).

### Ion imbalance, starch and proline content and their role in senescence

We found changes in ion content in seedlings (Kocourková *et al*., 2020) and leaves (this work) of plants treated with high Mg^2+^. At WT, the K^+^ content of plants treated with high Mg^2+^ was lower compared to untreated controls. Moreover, K^+^ content under high Mg^2+^ conditions was significantly lower in *pldα1* plants than in WT. Similarly, *pldα1* and WT plants also differed in P content under high-Mg^2+^ conditions; *pldα1* plants had lower P contents than WT. However, the P content of WT did not differ between control and high-Mg^2+^ conditions.

Potassium deficiency has been reported to induce leaf senescence in Arabidopsis and cotton. Interestingly, there is also strong evidence that jasmonic acid is involved in potassium deficiency - induced leaf senescence (Armengaud *et al*., 2004; Cao *et al*., 2006 ; Hu *et al*., 2016; Li *et al*., 2012). We demonstrated an increase in JA level after high Mg^2+^ treatment in both WT and *pldα1*, and that the accumulation of JA in *pldα1* was higher than that of WT. Further experiments are needed to determine whether potassium deficiency, JA accumulation, and leaf senescence are related.

Leaf starch is synthesised during the day and mobilised during the following night to provide a steady supply of carbon and energy. Starch also mediates plant responses to abiotic stresses such as water deficit, high salinity or extreme temperatures. Most studies considered that starch content in leaves decreases in response to abiotic stresses. However, there are also reports that starch accumulation increases in Arabidopsis under stress (Kaplan and Guy, 2004; Skirycz *et al*., 2009). In our work, we observed increased starch accumulation under high Mg^2+^ conditions in WT plants, and starch accumulation was even higher under these conditions in *pldα1* plants. The opposite effect of high Mg^2+^ was observed by Guo *et al*. (2014). The authors found lower leaf starch level in Arabidopsis WT under high Mg^2+^ conditions than under control conditions. It is not clear why such different results occurred. One possible explanation could be the use of different ecotypes and experimental conditions. Guo *et al*. (2014) used an ecotype (Landsberg erecta), a higher Mg^2+^ concentration (32 mM) and long-term high Mg^2+^ stress, while we observed starch in leaves on the third day after treating Arabidopsis plants of ecotype Columbia 0 with 15 mM Mg^2+^. The relationship between high starch accumulation and leaf senescence has also been described (Huang *et al*., 2018; Oda-Yamamizo *et al*., 2016; Schaffer *et al*., 1991; Xiao *et al*., 2020). A possible link between PLDα1 and altered starch accumulation could be the PA binding protein glyceraldehyde-3-phosphate dehydrogenase (GAPDH) (McLoughlin *et al*., 2013), as seedlings with genetically reduced GAPDH activity accumulated higher amounts of starch compared to WT (Yang *et al*., 2015). Moreover, phosphorus deficiency increases leaf sugars and starch content (Cakmak *et al*., 1994; Hermans *et al*., 2006) and we found lower phosphorus content in the leaves of *pldα1* plants compared to WT.

We observed proline accumulation in plants exposed to high Mg^2+^ conditions. Proline is a well-known molecule involved in adaptation to stress by e.g. balancing cellular redox potential, scavenging free radicals and stabilising subcellular structures (Kaur and Asthir, 2015; Szabados and Savoure, 2009). A relationship between proline metabolism and leaf senescence has been previously noted and discussed (Zhang and Becker, 2015). Proline content significantly increased in detached rice leaves during senescence (Wang *et al*., 1982). On the other hand, proline catabolism appears to be up-regulated in Arabidopsis during natural leaf senescence (Funck *et al*., 2010). Moreover, experiments with inhibition of PLD activity by 1-butanol during salt stress showed that PLD appears to be a negative regulator of the delta-1-pyrroline-5-carboxylate synthase 1 gene, which controls proline biosynthesis (Thiery *et al*., 2004). We have demonstrated that exposure of *pldα1* to high Mg^2+^ resulted in decreased proline accumulation compared to WT. Thus, PLDα1 appears to be a positive regulator of proline synthesis. Why PLDα1-depleted plants have less proline under high-Mg^2+^ conditions is unclear. The difference in proline content between WT and *pldα1* is significant only after prolonged exposure (4 days) to high Mg^2+^, and it is therefore possible that this is a side effect of earlier changes caused by high Mg^2+^ rather than a direct regulation of proline metabolism by PLDα1. However, it is possible that proline helps the plants to cope with the stress caused by high Mg^2+^, and the lower proline content of *pldα1* may contribute to the manifestation of senescence in plants with dysfunctional PLDα1. Interestingly, significant differences in proline content were observed between WT and phosphoenolpyruvate carboxylase 3 (PEPC3) knockout in Arabidopsis under control and salt stress conditions and PEPC3 was identified as the PA-binding protein (Testerink *et al*., 2004).

### Mechanism of PLDα1 involvement in leaf senescence

PLDα1 has been described to be involved in a variety of biological processes. PLDα1 knockout or antisense-suppressed plants show alterations in water loss, reactive oxygen species production (ROS), response to abscisic acid (ABA) and stomatal movement (Zhang *et al*., 2004; Zhang *et al*., 2009); Mishra, 2006 #9472}, salt stress (Bargmann *et al*., 2009; Yu *et al*., 2015; Yu *et al*., 2010), freezing sensitivity (Rajashekar *et al*., 2006) and seed aging (Devaiah *et al*., 2007). The involvement of PLDα1 in senescence has also been documented. In 1997, Fan *et al*. (Fan *et al*.) observed that treatment of detached leaves with abscisic acid and ethylene led to accelerated senescence and increased level of PLDα mRNA, protein and activity. Using the PLDα antisense construct, they then prepared plants with reduced PLDα1 expression. Suppression of PLDα had no effect on natural plant growth and development. Even in the absence of abscisic acid and ethylene, the detached leaves of the PLDα-deficient and WT plants showed similar rate of senescence.

However, the senescence rate of detached leaves of transgenic plants treated with ABA or ethylene was slower than that of detached leaves from WT. Later, Jia *et al*. (2013) showed that the application of n-butanol, an inhibitor of PLD, and N-acylethanolamine (NAE) 12:0, a specific inhibitor of PLDα1, delayed ABA-promoted senescence to different extents. Phospholipase Dδ (PLDδ) was also induced in detached leaves treated with ABA, and suppression of PLDδ delayed ABA and ethylene - promoted senescence in Arabidopsis (Jia and Li, 2015). These data suggest that suppression of both PLDα1 and PLDδ blocks membrane lipid degradation, which ultimately delays ABA-promoted senescence. Thus, PLDα1 and PLDδ appear to be important mediators that play a positive role in phytohormone-promoted senescence in detached leaves. However, in this work, we showed that PLDα1 likely serves as a negative regulator of senescence. We observed that PLDα1 activity and PLDα1 abundance increase during senescence triggered by high Mg^2+^ content. An increase in PLDα1 expression has also been described during age-related leaf senescence (Xiao *et al*., 2010). However, in our case, the increase in PLDα1 activity and PLDα1 abundance is probably not related to the increased membrane degradation described above, because *pldα1-1* plants exhibited significantly higher senescence compared with WT. Therefore, another regulatory mechanism by which PLDα1 is involved in the regulation of senescence induced by high Mg^2+^ levels must play a role.

In general, there are two molecular ways by which PLDα1 may regulate other events. The first is linked with PLDα1 activity which leads to the production of the second messenger phosphatidic acid and free head group, and the second is protein-protein interaction. In the case of PLDα1, both scenarios have been documented. A combination of both mechanisms is also possible and has been described in the case of PLDα1 involvement in ABA responses (see below). PLD-derived phosphatidic acid is produced in response to various biotic and abiotic stresses such as plant defence, wounding, salt, drought, cold, and heat stress (Yao and Xue, 2018). In salt stress induced PLDα1 produces PA, which stimulates mitogen-activated protein kinase 6 (MPK6) to regulate the MAPK cascade (Yu *et al*., 2010). PA also influences microtubule stabilisation by interacting with MAP65-1 during the Arabidopsis salt stress response (Zhang *et al*., 2012b). In addition, ABA, part of the salt stress response, activates PLDα1. PA subsequently binds to ABI1, a protein phosphatase 2C, a negative regulator of the ABA response. Binding of PA decreases the phosphatase activity of ABI1. Thus, PA, which is produced by PLDα1, inhibits the function of the negative regulator ABI1 and thereby promotes the signalling of ABA (Zhang *et al*., 2004). Since ABA functions as a positive regulator of leaf senescence, PLDα1 could also play the role of a positive regulator of leaf senescence. However, in our study, this is not the case because PLDα1 is a negative regulator of leaf senescence induced by high Mg^2+^ content. PLDα1 also plays a role in ABA-induced stomatal closure. It has been shown that PA, which was produced by PLDα1, binds to the NADPH oxidase RbohD (respiratory burst oxidase homolog D) and RbohF of Arabidopsis and that this binding stimulates NADPH oxidase activity and thus the production of ROS in guard cells (Zhang *et al*., 2009).

A number of PA-binding proteins have been found (Yao and Xue, 2018). CTR1 (CONSTITUTIVE TRIPLE RESPONSE1) is another example of PA binding protein (Testerink *et al*., 2004; Testerink *et al*., 2007). CTR1 is a Ser/Thr protein kinase that functions as a negative regulator of ethylene signalling. Loss of CTR1 function has been shown to promote the senescence process upon dark treatment, suggesting that CTR1 plays a role as a negative regulator of leaf senescence (Li *et al*., 2017). We did not measure ethylene levels in our experimental setup. However, ethylene is known to be an endogenous modulator of senescence, including leaf senescence. PLDα1 protein interaction, the second possible PLDα1 regulatory mechanism, is also involved in the regulation of ABA responses. PLDα1 interacts with components of heterotrimeric G protein signalling, GPA1 (Gα) and Gβ proteins (Gookin and Assmann, 2014; Zhao and Wang, 2004). The interaction of PLDα1 with GPA1 stimulates the GTPase activity of GPA1 (Zhao and Wang, 2004). PLDα1 also interacts with RGS1 protein (regulator of G protein signalling). RGS1 likely inhibits the GAP activity of PLDα1 (Choudhury and Pandey, 2016). To further impact the specificity of this pathway, PA, the product of PLDα1 activity, binds to RGS1 and inhibits its GAP activity. Interestingly, GPA1-, Gβ-as well as RGS1 knock-out plants showed altered salt stress-induced senescence (Colaneri *et al*., 2014).

In summary, high Mg^2+^ induces leaf senescence and many of the physiological changes associated with leaf senescence induced by high Mg^2+^ are under the control of PLDα1. Subsequent studies should elucidate the precise molecular mechanism of this PLDα1 control.

## Supplementary data

Fig. S1. PA amount increases in response to high-Mg^2+^ stress in wt but not in *pldα1-1*.

Fig. S2. Phospholipase Dα1 amount and activity increases in response to high Mg^2+^ stress in youngest leaves

Table S1: Hormone analysis

Table S2: List of primers

## Acknowledgements

This research was supported by the Czech Science Foundation (grant No. 17-00522S) and the Ministry of Education, Youth and Sports of the Czech Republic (European Regional Development Fund-Project ‘Centre for Experimental Plant Biology’ no. CZ.02.1.01/0.0/0.0/16_019/0000738). The authors wish to thank Kateřina Vltavská for excellent technical assistance.

## Author Contributions

DK and JM designed the study and wrote the manuscript. DK, KK, TP, and MD performed the experiments. All authors reviewed and edited the manuscript.

